# Vaccination of cattle with a virus-vectored vaccine against a major membrane protein of *Mycobacterium a*. subsp. *paratuberculosis* elicits CD8 cytotoxic T cells that kill intracellular bacteria

**DOI:** 10.1101/2023.11.20.567939

**Authors:** Asmaa H. Mahmoud, Gaber S. Abdellrazeq, Valentina Franceschi, David A. Schneider, John P. Bannantine, Lindsay M. Fry, Victoria Hulubei, Giovanna De Matteis, Kun Taek Park, William C. Davis, Gaetano Donofrio

## Abstract

Infection of cattle with *Mycobacterium a*. subsp. *Paratuberculosis (Map)*, the causative agent of paratuberculosis, induces an immune response directed toward a 35 kD major membrane protein (MMP) of *Map*. CD8 cytotoxic T cells (CTL) are elicited when peripheral blood mononuclear cells from healthy cattle are incubated ex vivo with antigen-presenting cells (APC) primed with bacterial recombinant MMP. Ex vivo development of CTL was MHC-restricted and required the presence of both CD4 and CD8 T cells. The gene *MAP2121c,* encoding MMP, was modified to express a modified form of MMP (p35NN) in a mammalian cell line, also capable of eliciting an ex vivo CTL response. In the present study, the modified gene for p35NN was placed into a BoHV4 vector to determine the potential use of BoHV-4AΔTK-p35NN as a peptide-based vaccine. Subcutaneous vaccination of healthy cattle with BoHV-4AΔTK-p35NN elicited a CTL recall response, as detected ex vivo. Further studies are warranted to conduct a challenge study to determine if CD8 CTL elicited by vaccination with BoHV-4AΔTK-p35NN prevents the establishment of a persistent infection by *Map*.

## 1. Introduction

*Mycobacterium a*. subsp. *paratuberculosis* (*Map*) is one of multiple lineages of mycobacteria that have evolved over the millennia, that have acquired the ability to infect and cause disease in multiple species, including humans (1) (reviewed in Bachmann et al. (2)). It is one of the major mycobacterial pathogens impacting the livestock industry and human health. Like other mycobacterial pathogens, it has been difficult to control. Initial infection induces an immune response that can control but not clear the pathogen, which results in a persistent infection. The lack of understanding of how a pathogen can establish a persistent infection in the presence of an immune response has, until now, impeded progress in developing a vaccine for *Map* and other mycobacterial pathogens. However, a seminal observation made by investigators studying the role of the stringent response in survival of *Mycobacterium tuberculosis* (*Mtb*) in a mouse model, discovered that the gene *rel*, regulator of the stringent response, is essential for survival in the mammalian host (3). When the gene is present, *Mtb* can establish a persistent infection. When the gene is deleted, the mutant *Mtb* can only establish a transient infection despite the initial formation of characteristic granulomas. The potential importance of this finding to other mycobacterial infections was revealed when the same effect on persistent infection was observed with a *rel* deletion mutant of *Mycobacterium a. paratuberculosis* (*Maprel*) in cattle and goats, natural hosts of *Map* (4, 5).

Analysis of the recall response ex vivo revealed vaccination with the mutant elicited the development of CD8 cytotoxic T cells (CTL) that could kill intracellular *Map* (6). Further analysis demonstrated one of the targets of the response is a 35 kD major membrane protein (MMP) (6). Comparison of the recall responses elicited with APC primed with *Maprel* or with MMP demonstrated both methods elicited comparable proliferative responses of CD4 and CD8 T cells, and development of CD8 T cells with similar CTL activity. No proliferative or CTL activity was detected in natural killer cells (NK) or γδ T cells also present in the preparations of PBMC.

Subsequent studies revealed ex vivo stimulation of PBMC with MMP elicited primary immune responses in CD4 and CD8 T cells with CD8 T cells acquiring CTL activity (6). This suggested it might be possible to develop a MMP-based vaccine against *Map*. The gene encoding MMP, *MAP2121c*, was modified for expression in mammalian cells to explore this possibility. The modified gene was placed in an expression cassette to produce modified MMP for analysis. Stimulation of PBMC with APC primed with a tPA-MMP-2mut expressed peptide, elicited CD4 and CD8 T cell proliferative responses, and CTL activity in CD8 T cells that was comparable to that elicited by the *Map* deletion mutant and MMP expressed in *E. coli* (eMMP).

As reported in the present study, the synthetic ORF tPA-MMP-2mut was used to develop and test a virus vectored vaccine using the expression vector, BoHV-4AΔTK, developed by Donofrio et al. (7). The results show the modified MMP is expressed and elicits development of CD8 CTL, as detected by analysis of the recall response ex vivo with APC primed with either *Maprel* or eMMP.

## 2. Materials and methods

### 2.1 Generation of constructs to develop a virus vectored MMP candidate vaccine

The synthetic ORF tPA-MMP-2mut, now called p35NN, was previously described (8). In this synthetic ORF, the Kozak consensus sequence and human tissue plasminogen activator signal peptide (tPA) was added to the 5’ terminus, and an AU1 peptide tag was added to the 3’ terminus. In addition, the first two predicted N-glycosylation sites were mutated to generate p35NN. The primer pair NheI-p35-sense (5’-CCCCGCTAGCCCACCATGGACGCTATGAAGAGGGGCCTGTGCTGC-3’) and SmaI-p35-antisense (5’-CCCCCCCGGGTTAGATGTACCGGTAGGTGTCCTTGTACTC-3’) was used to insert a NheI restriction site at the 5’ end and a SmaI restriction site at the 3’end of the ORF. The NheI/SmaI cut amplicon was cloned into the shuttle vector pINT2EGFP (9) and cut with the same enzymes, generating pTK-CMV-p35NN-TK; thereby placing the p35NN ORF under the transcriptional control of the immediate early gene promoter of human cytomegalovirus (CMV), followed by the bovine growth hormone polyadenylation signal (pA) and flanked by BoHV-4 TK sequences.

### 2.2 Transient transfection and secretion of p35NN from HEK 293T cells

Cells were seeded at 3x10^5^ cells/well in 6-well plates and incubated overnight at 37°C and 5% CO_2_ in a humidified incubator. Cells were then incubated for 6 hours with a transfection mix containing 3 µg plasmid DNA and PEI (ratio 1:2.5 DNA-PEI) in Dulbecco’s Modified Essential Medium (DMEM) high glucose (Euroclone) without serum. After incubation, the transfection mix was replaced by fresh complete Eagle’s Modified Essential Medium (cEMEM: 100 IU/mL of penicillin, 100 μg/mL of streptomycin, 0.25 μg/mL of amphotericin B, 1 mM of sodium pyruvate and 2 mM of L-glutamine), with 10% Fetal Bovine Serum (FBS), and incubated for 24 hours at 37°C and 5% CO_2_ in a humidified incubator. To test supernatant protein expression, the transfection solution was replaced with fresh DMEM/F12 (ratio 1:1) medium without FBS and incubated for 72 hours at 37°C and5% CO_2_ in a humidified incubator. Cell supernatant was then collected and analyzed by immunoblot. Cell supernatants, obtained from HEK 293T cells transfected pTK-CMV-p35NN-TK or pEGFP-C1, were collected after 72 hours in serum free medium DMEM-F12 secretion condition and analyzed through 10% SDS–PAGE gel electrophoresis. Proteins were then transferred to PVDF membranes by electroblotting, and membranes were incubated with anti-AU1 rabbit polyclonal antibody (A190-125A, Bethyl laboratories Inc.) diluted 1:10,000, washed, and then incubated with a goat anti-rabbit secondary antibody labelled with horse radish peroxidase (Sigma), diluted 1:15,000 and visualized by enhanced chemiluminescence (Clarity Max western ECL substrate, Bio-Rad).

### 2.3 Bacterial Artificial Chromosome (BAC) Recombineering and Selection

The PvuI linearized pTK-CMV-p35NN-TK expression cassettes was used for heat-inducible homologous recombination in SW102 *E. coli*, containing the BAC-BoHV-4-A-TK-KanaGalK-TK genome and targeted into the TK locus with KanaGalK selector cassette. After recombineering, only those colonies that were kanamycin resistant and chloramphenicol positive were isolated and grown overnight in 5 mL of LB (Luria-Bertani) liquid medium broth containing 12.5 mg/mL of chloramphenicol. BAC-DNA was purified and analyzed through HindIII restriction enzyme digestion. DNA was separated by electrophoresis in a 1% agarose gel. Original detailed protocols for recombineering can also be found at the recombineering website (https://redrecombineering.ncifcrf.gov/). Non-isotopic Southern blotting was performed with a probe specific for the p35NN sequence. DNA from 1% agarose gel was capillary transferred to a positively charged nylon membrane (Roche) and cross-linked by UV irradiation by standard procedures. The membrane was pre-hybridized in 50 mL of hybridization solution (7% SDS, 0.5 M phosphate, pH 7.2) for 1 hour at 65°C in a rotating hybridization oven. P35NN probe labeled with digoxigenin was generated by PCR with primers described above. The PCR amplification reaction was carried out in a final volume of 50 µL, containing 10 mM Tris–hydrochloride pH 8.3, 5% dimethyl sulfoxide (DMSO), 0.2 mM deoxynucleotide triphosphates, 2.5 mM MgSO_4_, 50 mM KCl, 0.02 mM alkaline labile digoxigenin-dUTP (Roche) and 0.25 µM of each primer. 100 ng of DNA was amplified over 35 cycles, as follows: 1 min denaturation at 94°C, 1 min annealing at 60°C, and 2 minutes elongation with 1U of Taq DNA polymerase (Thermo Fisher Scientific) in addition to 1 µl of Digoxigenin-11-dUTP, alkali-labile (Roche) at 72°C.

### 2.4 Cell Culture Electroporation and Recombinant Virus Reconstitution

BEK or BEK*cre* cells were maintained as a monolayer with cEMEM + 10% FBS. When cells were sub-confluent (70-90%) they were split to a fresh culture flask and incubated at 37°C, 5% CO_2_. BAC-DNA (∼5 µg) was electroporated in 600 µL DMEM without serum (Bio-Rad Gene pulser Xcell, 270 V, 1500 µF, 4-mm gap cuvettes) into BEK and BEK*cre* cells. Cells were left to grow until the appearance of syncytia of infected cells (CPE).

### 2.5 Virus and Viral amplification

BoHV-4-A-CMV-p35NNΔTK was propagated by infecting confluent monolayers of BEK cells at a multiplicity of infection (MOI) of 0.5 tissue culture infectious doses 50 (TCID_50_) per cell and maintained in medium with only 2% FBS for 2 hours. Medium was replaced with fresh cEMEM with 10% FBS. When the majority of the cell monolayer displayed the CPE (∼72 hours post infection), the virus was harvested by freezing and thawing cells three times and pelleting the virions through a 30% sucrose cushion. Virus pellets were then resuspended in cold DMEM without FBS. The tissue culture infectious dose (TCID_50_) was determined on BEK or MDBK cells by limiting dilution.

### 2.6 Virus Growth Curves

BEK cells were infected with BoHV-4-A-CMV-p35NNΔTK and BoHV-4-A at a MOI of 0.1 and incubated at 37°C for 4 hours. Infected cells were washed with serum-free cEMEM and then overlaid with cEMEM. The supernatants of infected cultures were harvested daily, and the amount of infectious virus was determined by limiting dilution on BEK cells. Viral titer differences between each time point are the average of triplicate measurements ± standard errors of the mean (*P* > 0.05 for all time points as measured by Student’s t-test).

### 2.7 Animals

A total of ten naïve Holstein steers were obtained from the Knott Dairy, a *Map*-free facility at Washington State University (WSU). All animal care and experimental procedures were conducted in compliance with the protocols approved by the Institutional Animal Care and Use Committee, WSU (ASAF 6542). Animals were grouped and treated as follows: One group of four steers was immunized with a subcutaneous (SC) injection of 10^7^ TCID_50_ of the virus vectored MMP (BoHV-4-A-CMV-p35NNΔTK). The second group of four steers was inoculated SC with 10^7^ TCID_50_ of the vector containing eGFP (BoHV-4-A-eGFPΔTK). The third group of two age-matched (1 yr) steers were left without inoculation. The steers were kept in an open feedlot and maintained by the WSU animal care staff and used as a source of blood throughout the duration of the study.

### 2.8 Preparation and culture of bacteria

The *Maprel* deletion mutant used in this study was developed from *Map* K-10 strain (*Map_K10_*) using a site-directed allelic exchange, as previously described (10). The *Map_K10_* and *Maprel* were grown as described in (6). When needed, master stocks were prepared and the optical density at 600 nm (OD_600_) was used to estimate the final bacterial number (10). Bacteria were diluted to a multiplicity of infection (MOI) needed to conduct each experiment as described below.

### 2.9 Cell separation and culture

Blood (∼ 200 mL) was collected from steers at two- and six-weeks postvaccination to analyze recall responses of isolated peripheral blood mononuclear cells (PBMC) as previously described (6). In brief, PBMC were suspended in a complete culture medium with and without antibiotics (cRPMI-1640) and then distributed in 6-well culture plates (2 × 10^6^/mL in 5 mL of RPMI/well) in duplicate to achieve two technical replicates. Live *Maprel* (1 × 10^6^/mL/well, MOI 0.5:1) or *E. coli* expressed MMP (eMMP, 5 G/ml) (11) were added to each of the two PBMC wells respectively. Two wells of PBMC were left unstimulated (mock) as negative controls. Two identical sets of PBMC cultures were prepared for analysis of the recall response; one set to analyze the proliferative response by flow cytometry (FC) and one to analyze the killing of intracellular bacteria by CD8 T cells by qPCR.

One portion of PBMC was used to generate monocyte-derived macrophages (MoMac) for use as target cells in the killing assay as described in (12).

### 2.10 Analysis of the recall response

#### 2.10.1 Flow cytometric analysis of the memory recall response to *Maprel* and eMMP stimulated PBMC

PBMC were cultured for 6 days in presence of either *Maprel* or eMMP or left unstimulated (mock). The cells were washed and labeled for the FC analysis as described in (6). In brief, after the cells were washed twice with first wash buffer (FWB: PBS containing 0.01% w/v sodium azide, 0.02% v/v horse serum, and 10% v/v acid citrate dextrose) the cells were incubated with mAbs listed in Table 1 (1 μg of mAb/10^6^ cells) for 15 min in the dark on ice. Cells were then washed twice using FWB and re-suspended in 100 μL of fluorochrome conjugated goat anti-mouse isotype-specific secondary mAbs. Specific isotype controls were used in the experiment. Cells were incubated for 15 min in the dark on ice, then washed twice with a second wash buffer SWB, as FWB but without horse serum. After the final wash, cells were re-suspended in 2% PBS-buffered formaldehyde and stored at 4°C until examined by FC. Data were acquired with a BD FACS Calibur flow cytometer (BD, Immunocytometry Systems, USA). Approximately 5 × 10^5^ events were collected for each sample. A sequential gating strategy was used to isolate CD4 and CD8 T cells for analysis as illustrated in Fig. 1A. A side light scatter (SSC) vs forward light scatter (FSC) gate was used to isolate lymphocytes for analysis. The proportions of activated memory CD45R0 CD4 and CD45R0 CD8 T cells in the total CD4 or CD8T-cell pools were determined by using electronic gates as shown in Fig. 1B. FCS Express software (De Novo Software, Pasadena, CA) was used to analyze all FC data.

**Fig. 1.**
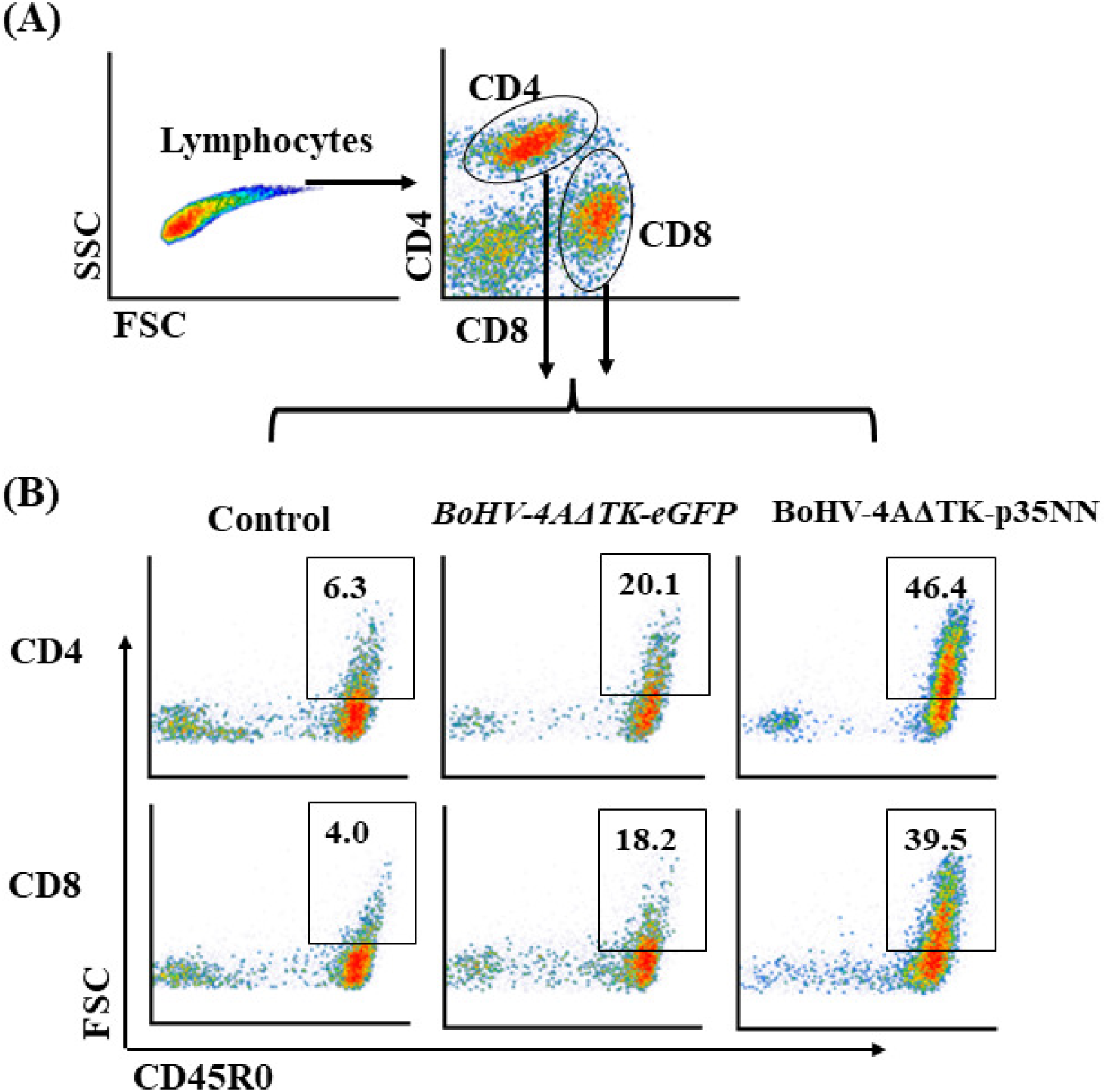
Flow cytometric profiles illustrating gating strategy used to obtain data. A) First gate placed on lymphocytes. B) Selective gates were placed on CD4 and CD8 T cells labeled with mAbs specific for CD4, CD8, and CD45R0 conjugated with different fluorochromes. The labeled CD4 and CD8 are displayed in forward light scatter (FSC) vs CD45R0 to visualize memory cell response to eGPF and BoHV-4-p35MMΔTK.

**Table 1.**
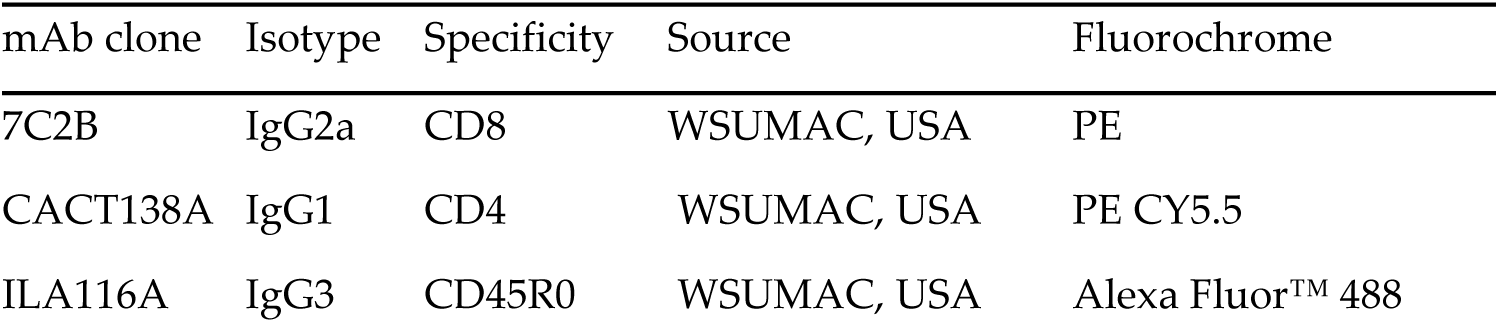
List of monoclonal antibodies used in this study.

#### 2.10.2 Intracellular killing assay for determination the cytotoxic activities of *Maprel* and eMMP-stimulated PBMC

The intracellular killing assay was used as previously described (6) to determine the proportion of bacteria killed by CD8 CTL generated in the recall response to BoHV-4-A-CMV-p35NNΔTK in comparison to the eGFP containing vector and unvaccinated controls. PBMC stimulated 6 days with *Maprel* or eMMP were collected and cocultured for an additional 24 hrs with MoMac target cells infected with live *Map_k10_* at MOI of 10:1 (2 × 10^7^ *Map_k10_* to ∼2 × 10^6^ MoMac/well). On the following day, coculture wells were collected individually and permeabilized with 1× saponin solution (0.5% w/v in PBS 10×, Thermo Fisher Scientific, CA, USA) for 15 minutes at 37°C in a humidified incubator. A set of controls (100% live, 50% live/50% killed, and 100% killed) were prepared from known mixtures of live and dead *Map_k10_* as described in (6) to cover the dynamic range of bacteria live/dead variable proportions in the infected target cells. All saponin treated cultures were washed twice in DNase-free dH_2_O, treated with a photoreactive propidium monoazide dye (PMA, Biotium, Fremont, CA, USA) in a final dye concentration of 50 μM, and then DNA was extracted (6). A preparation of *Map_k10_* grown to log phase (4 × 10^7^) was included during DNA extraction to obtain pure *Map_k10_* gDNA for use to generate a standard curve. The single-copy *F57* gene is specific for *Map* and was used in the TaqMan qRT-PCR to determine the proportions of live *Map_k10_* within each culture as described by Kralik et al. (13) and Abdellrazeq et al.(6). All PCR reagents including primers, TaqMan probe, and TaqMan universal master mix were supplied by Applied Biosystems (Thermo Fisher Scientific, CA). The qRT-PCR reaction and cycling parameters were performed according to Schönenbrücher et al. (14) using a StepOnePlus Real-Time PCR System machine (Applied Biosystems, CA). Each sample was run in triplicate. The mean cycle threshold (C_T_) values of the 3 replicates were determined using StepOne Software v2.1 (Applied Biosystems, CA) and used to calculate the relative proportion of viable bacteria in each sample. Data were considered valid if the following criteria were met: PCR efficiency between 90 and 110%, regression coefficient (R^2^) > 0.99, and standard deviation ≤ 0.250.

### 2.11 Statistical analysis

A generalized linear mixed model analysis was conducted on each dataset using the SAS software procedure PROC GLIMMIX (SAS Institute Inc., Cary, NC, USA). Each model included the main effects terms for vaccination group (none, eGFP, p35NN) and ex vivo antigen stimulus (mock, eMMP, *Maprel*), the interaction term, and a random intercept for the subjects (steers nested within vaccination groups). Separate analyses were conducted for each T cell type (CD4 or CD8) at postvaccination weeks two and six. The memory recall response measured as proportion of activated (CD45R0) T cells was modeled as a beta distribution whereas the intracellular bacterial killing response (C_T_ value) was best modeled as a Gaussian distribution. Specific linear contrasts were constructed to compare responses to ex vivo mock stimulation (baseline) between vaccination groups. Similarly, specific linear contrasts were constructed to compare responses to each ex vivo antigen stimulus (eMMP and *Maprel*) to mock stimulation within each vaccination group. The improved method of Kenward and Roger (DDFM=KR2) was used to compute denominator degrees of freedom. Significance values were adjusted for multiplicity using the technique of Holm (step-down Bonferroni; *P_Holm_* < 0.05). The modeled responses are summarized in figures as the predicted least squares means and 95% confidence limits.

## 3. Results

### 3.1 Development of a recombinant BoHV-4 expressing p35NN

A recombinant BoHV-4 virus expressing a secreted mutant form of p35 (tPA-MMP-2mut) was used in the present study (8) to develop an expression cassette, CMV-p35NN. Expression was assessed by transient transfection in HEK 293T cells. Transfected cells secreted p35NN in the supernatant as detected by western immunoblotting (Fig. 2A). A recombinant BoHV-4 delivering CMV-p35NN, BoHV-4-A-CMV-p35NNΔTK, was generated starting with the genome of an apathogenic BoHV-4 strain (BoHV-4-A) (15) cloned as a BAC. The TK BoHV-4-A genome locus was used as the integration site for the CMV-p35NN expression cassette. The BoHV-4 TK genomic region is strongly conserved among BoHV-4 isolates (16), ensuring the stability of the genomic locus from potential recombination. The TK locus does not interfere with viral replication in vitro and heterologous protein expression is maintained (9, 17) (9, 17–21) (22–24). Restriction enzyme linearized pTK-CMV-p35NN-TK was used for heat-inducible homologous recombination in SW102 *E*. *coli* containing pBAC-BoHV-4-A-KanaGalKΔTK (15.1016/j.vaccine.2008.09.023 ) (Fig. 2B) to generate pBAC-BoHV-4-A-CMV-p35NNΔTK. Selected clones were first analyzed by *HindIII* restriction enzyme digestion and Southern blotting (Fig. 2B, C). Because heat-inducible recombination in SW102 *E*. *coli* and repeated passages could establish altered bacterial phenotypes, due to aberrant recombinase transcription, SW102 *E*. *coli* carrying pBAC-BoHV-4-A-CMV-p35NNΔTK were serially cultured for over 20 passages and checked by *HindIII* restriction enzyme digestion. No differences were detected among restriction patterns at various passages (data not shown), thus ensuring the stability of the clones. To reconstitute infectious BAC-BoHV-4-A-CMV-p35NNΔTK, pBAC-BoHV-4-A-CMV-p35NNΔTK DNA was electroporated into BEK and BEK*cre* cells (15). The recombinant viruses reconstituted from electroporated BEK*cre* resulted in depletion of the BAC plasmid backbone containing the GFP expression cassette, as shown by the loss of GFP expression (Fig. 2D). Next, the growth characteristics of BoHV-4-A-CMV-p35NNΔTK were compared with that of the parent virus, BoHV-4-A. BoHV-4-A-CMV-p35NNΔTK demonstrated slightly slower replication kinetics compared to BoHV-4-A (Fig. 2E). Transgene expression was detected in the supernatant of BoHV-4-A-CMV-p35NNΔTK infected cells (Fig. 2F).

**Fig. 2.**
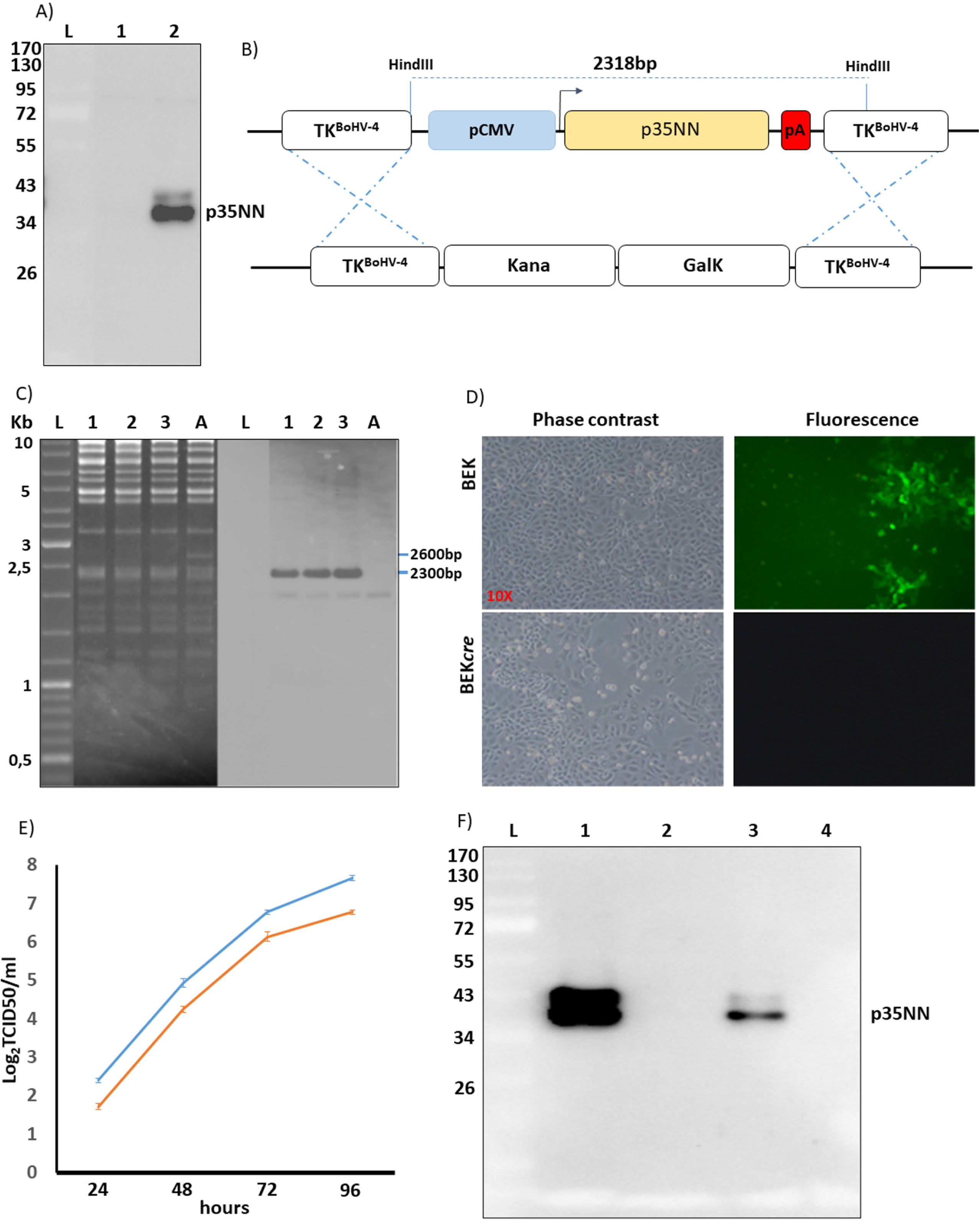
Summary characterization of BoHV-4-p35MMΔTK constructs and**. A)** Immunoblotting analyses conducted on supernatant from pEGFP-C1 (1) and pTK-CMV-p35NN-TK transfected HEK 293T cells **B)** Diagram (not to scale) illustrating the re-targeting event (i.e., replacement of the Kana/GalK cassette with the CMV-p35NN cassette) obtained by heat-inducible homologous recombination in SW102 *E.coli* cells containing pBAC-BoHV-4-A-TK-KanaGalK-TK. **C)** Representative, 2-deoxy-galactose resistant colonies, tested by *Hind* III restriction enzyme analysis and southern blotting performed with a specific probe for p35NN ORF. The 2,650 bp band corresponding to the non-retargeted pBAC-BoHV-4-A-TK-KanaGalK-TK control (lane A) is replaced by 2,300 bp band in pBAC-BoHV-4-p35NNΔTK (lanes 1, 2 and 3). **D)** Phase contrast and fluorescent microscopy images of the plaques formed by viable, reconstituted recombinant BoHV-4-p35NNΔTK after electroporation of the corresponding BAC DNA clones into BEK or *BEKcre* cells (magnification, ×10). **E)** Replication kinetics of BoHV-4-A and BoHV-4-p35NNΔTK . **F)** Immunoblotting analyses conducted on supernatant from BBMC cells (lane 1 and 2) and MDBK cells (lanes 3 and 4) infected with BoHV-4-tPA-MMP-2mutΔTK (lanes 1 and 3) or BoHV-4-A parental virus (lanes 2 and 4). Each lane was loaded with 15 µls of supernatant incubated for 72 hours post infection.

### 3.2 Recall response to vaccination with BoHV-4-A-CMV-p35NNΔTK

Preliminary studies revealed the virus vectored MMP could not be used ex vivo to prime APC to elicit a recall response. Therefore, eMMP and *Maprel* were used for eliciting a recall response. Although initial studies with the gene encoding MMP, modified for expression in a virus vector, indicated the modifications did not alter immunogenicity, it was not known if the epitopes expressed by the modified MMP were the same as those expressed in eMMP. It was also not known if the retained epitopes would elicit a recall response to the epitopes present in native MMP. Previous studies with eMMP had provided data indicating epitopes processed and presented concurrently by MHC I and MHC II were essential for eliciting a proliferative CTL response (25). *Maprel* was included to determine if vectored MMP used to vaccinate steers elicit a recall response to epitopes conserved on eMMP and native MMP expressed by *Maprel*.

### 3.3 Flow cytometric analysis of the recall response

The proportions of CD4 T and CD8 T cells expressing CD45R0 after six days culture of PBMC in the absence (mock) or presence of eMMP or *Maprel* are summarized in Figs. 1 and 3. The flow cytometric profiles in Fig. 1 illustrate how numerical data of memory CD4 and CD8 T cells were obtained. Fig. 3 provides a summary of the data. The baseline proportions of memory CD4 and CD8 T cells that expressed CD45R0 after PBMC from unvaccinated steers were cultured alone or in the presence of eMMP or *Maprel* were minimal but, relative to unvaccinated steers, significantly increased in comparison with both groups of vaccinated steers at two and six weeks post-vaccination (Fig. 3, # indicates 0.001 ≤ *P_Holm_* < 0.05 for differences in baseline proportions between vaccination groups). In contrast, robust increases from baseline were observed at both time points when PBMC from steers vaccinated with virus-vectored p35NN were cultured with eMMP or *Maprel* (Fig. 3, ** and *** respectively indicate *P_Holm_* < 0.001 and < 0.0001).

**Fig. 3.**
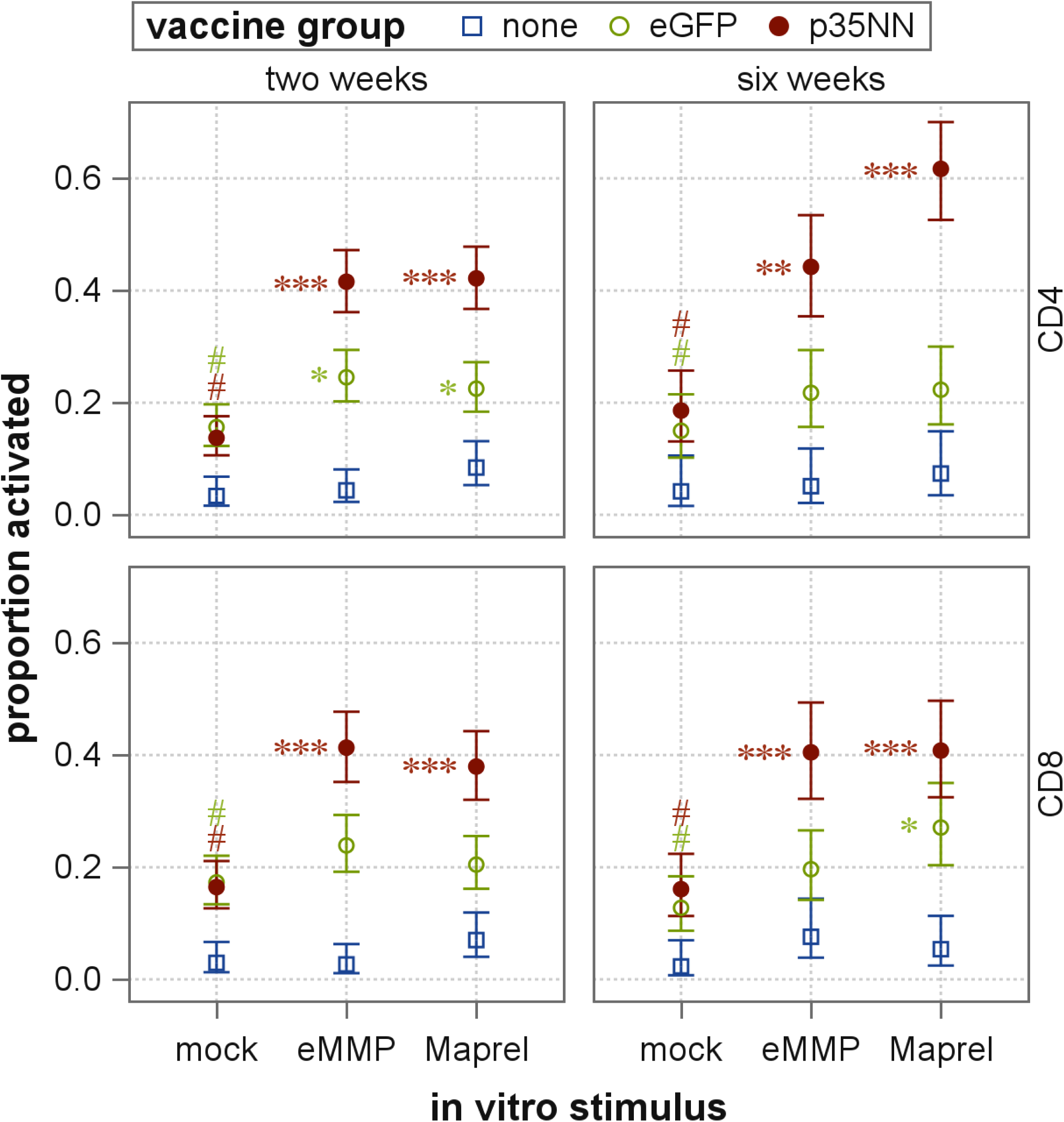
Comparison of the proliferative recall response to vaccination of steers with virus-vectored p35NN and virus-vectored eGFP at two and six weeks postvaccination. Only p35NN elicited a consistent proliferative recall response.

### 3.4 Analysis of the cytotoxic T cell recall response

Figure 4 shows an example of the *F57* gene C_T_ values measured for PBMC collected from a steer 2 and 6 weeks after vaccination with the virus-vectored p35NN. A summary of the *F57* gene C_T_ results for all vaccination groups and culture conditions at each time point is shown in Figure 5. There were no significant differences between vaccination groups in the baseline measures of C_T_ values (Fig. 5, mock). PBMC from unvaccinated steers cultured in the presence of eMMP or *Maprel* did not significantly change C_T_ values from baseline at either time point. For steers vaccinated with virus-vectored eGFP, culture in the presence of eMMP or *Maprel* did result in increased C_T_ values for PBMC collected at two-weeks post vaccination, but which were not sustained in PBMC collected at six-weeks (Fig. 5, * indicates 0.001 ≤ *P_Holm_* < 0.05 for differences relative to the corresponding baseline value). For steers vaccinated with virus-vectored p35NN, C_T_ values were increased when PBMC collected at two-weeks post-vaccination were cultured in the presence of eMMP or *Maprel*, which was sustained in PBMC collected at six-weeks only in response to eMMP. (Fig. 5, * and ** respectively indicate *P_Holm_* < 0.05 and < 0.001 for differences relative to the corresponding baseline value).

**Fig. 4.**
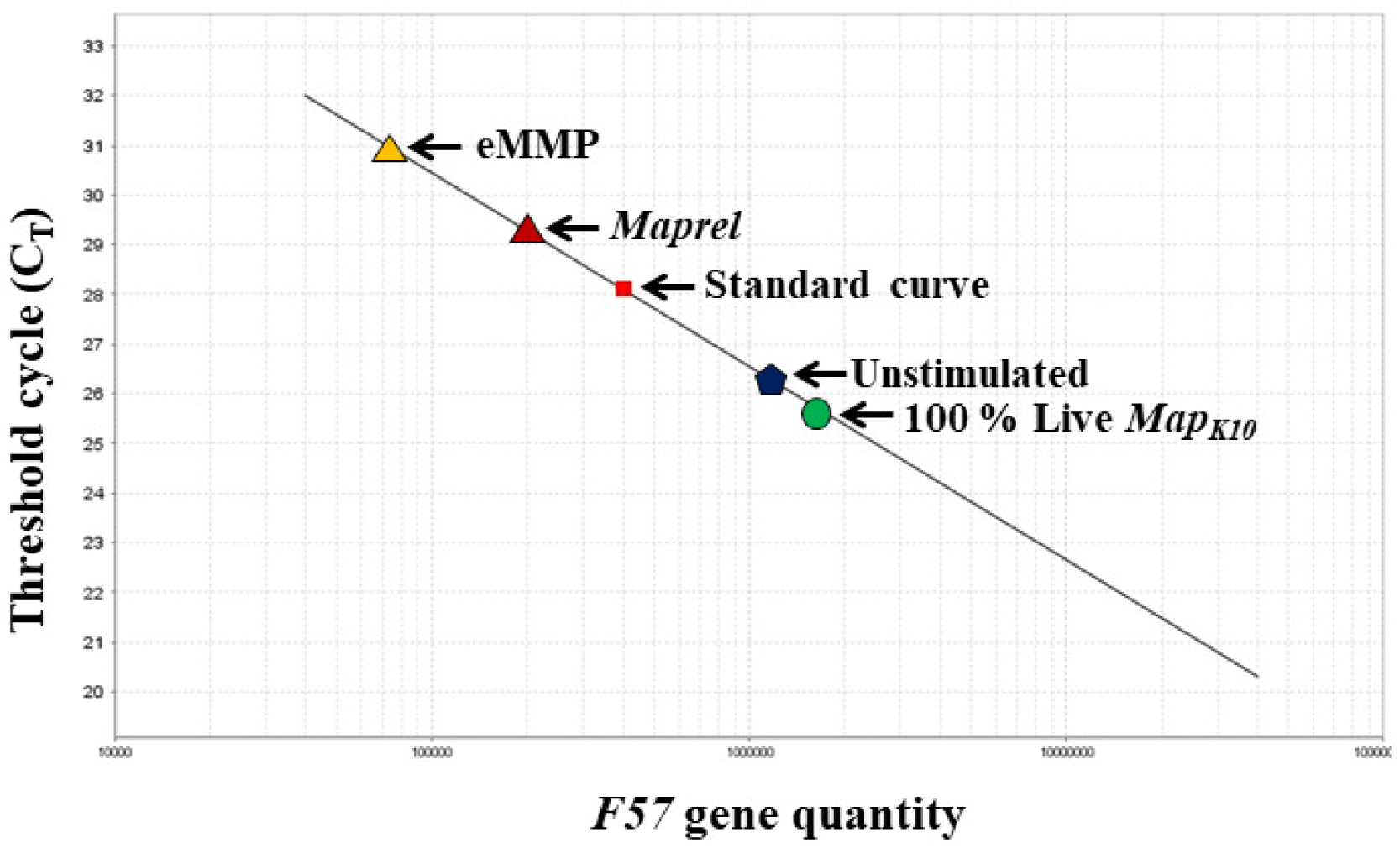
Illustration of a quantitative PCR curve set up with serial dilutions of DNA from live *Map*. DNA standards prepared from a single copy gene, F57, present in 100% live, 50% live, and 100% dead are placed on the DNA curve to plot the DNA dynamic range for quantifying the number of live bacteria remaining after incubation of infected target cells with memory CD8 T cells primed with vectored p35NN or controls.

**Figure 5.**
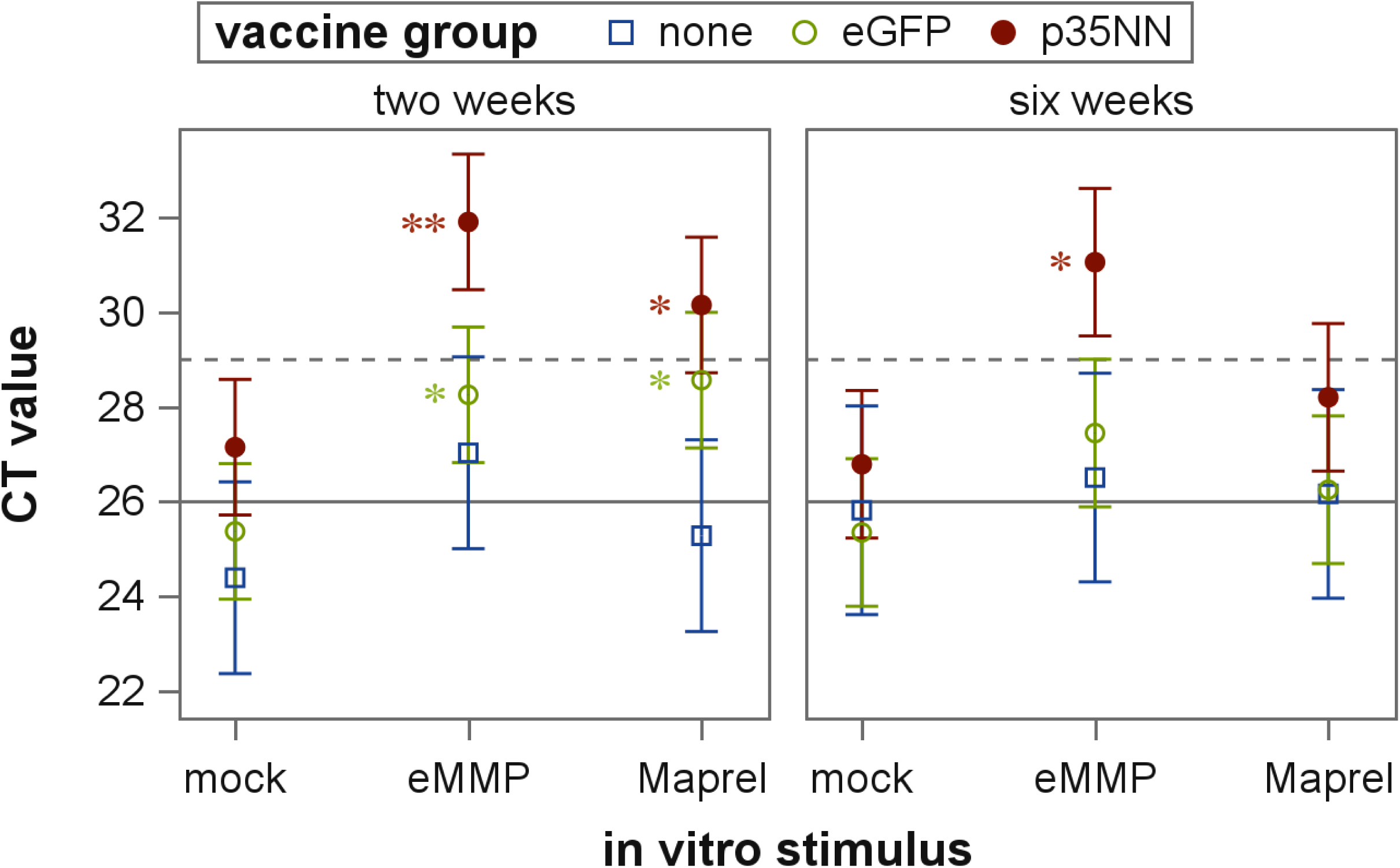
Comparison of live *Map* remaining in macrophage target cells after co-culture with CD8 CTL from steers vaccinated with virus-vectored p35NN or virus-vectored eGFP controls. Vaccination of steers with virus-vectored p35NN, but not virus-vectored eGFP, resulted in consistent killing of intracellular bacteria.

## 4. Discussion

The development of methods to use cattle as a model species to study the immune response to mycobacterial pathogens has afforded an opportunity to follow up on observations made with a mouse model which showed that deletion of *rel* abrogates the ability of *Mtb* to establish a persistent infection (26, 27). Similarly, a *Maprel* deletion mutant could not establish a persistent infection in cattle or goats (5). Furthermore, ex vivo studies with *rel* deletion mutants in *Map* and BCG demonstrated immediate CD4 and CD8 T cell responses were elicited with APC primed with the mutants, and CD8 CTL developed the ability to kill intracellular bacteria (6, 28). The 35 kD membrane protein, MMP, was shown to be one of the targets of in the *Maprel* mutant (6).

A comparable immune response is also obtained when APCs are primed with either eMMP or *Maprel* (6). Stimulation of naïve PBMC ex vivo with APC primed with eMMP elicited a CD8 CTL response. This response required simultaneous recognition of eMMP epitopes (presented in context MHC I and II molecules) to elicit primary and recall CTL responses (25). These findings indicate any peptide considered for developing a peptide-based vaccine must contain epitopes that will be presented in context of MHC I and II molecules (6). In all, the findings indicate the eMMP protein fulfilled the requirements for developing a peptide-based vaccine. The epitopes recognized by MHC I and MHC II were retained in the modified MMP expressed by the virus vectored p35NN.

An early attempt to develop a nano particle-based vaccine incorporating eMMP was not successful (29). Initial studies demonstrated eMMP could be encapsulated in a nano particle lipid carrier and elicit a recall response. However, follow up studies revealed it was not possible to develop a stable eMMP nanoparticle based vaccine. In the present study, we turned to a virus-vectored vaccine approach. The question posed was whether modification of the gene encoding eMMP for expression in a mammalian cell would retain MHC I and II restricted epitopes required for eliciting CTL. Modification of the gene was required to incorporate it into a shuttle vector, but the expressed modified-gene products still elicited a CTL response comparable to eMMP. Further modification and incorporation of the gene encoding MMP into the BoHV-4 vector was successful (BoHV-4-A-CMV-p35NNΔTK) and resulted in a post-vaccination response in steers that included a CTL response within two weeks. Further studies are planned to determine whether the immune response elicited by BoHV-4-A-CMV-p35NNΔTK is sufficient to block establishment of a persistent infection by *Map*. Similar studies are also planned to determine if the broader immune response elicited by the *Maprel* mutant elicits an immune response with similar or different protective activity.

## Conflicts of Interest

The authors declare that they have no competing interests.

## Author Contributions

AHM, GSA, LMF, JPB, and WCD conceived the study. KTP developed the *rel* deletion mutants used to conduct the studies. GD developed the MMP BoHV-4 vectored vaccine, AHM, GSA, and WCD participated in the design of the protocol to conduct the studies. AHM and GSA conducted the studies. AHM, GSA and DAS participated in statistical analysis of the data. AHM, GSA, WCD, LMF, VF, GDM, VH, KTP, JPD, GD, and DAS participated in the writing and interpretation of the results. WCD obtained the funding for the project. WCD oversaw and participated in all aspects of the study. All authors read and approved the final manuscript.

## Acknowledgments

The authors wish to acknowledge the excellent technical support and animal care provided by Emma Karol and her staff. WCD would like to recognize Sidney Raffel, Standford University School of Medicine, mentor and early pioneer studying mechanisms of pathogenesis of mycobacterial pathogens (1911–2013).

## Funding

This work was supported in part by the USDA National Institute of Food and Agriculture-Agriculture and Food Research Initiative Competitive Grant 2018–67015-28744 (W. C. Davis) and 1020620. Mention of trade names, proprietary products, or specified equipment do not constitute a guarantee or warranty by the USDA and does not imply approval to the exclusion of other products that may be suitable. USDA is an Equal Opportunity Employer. This study was also supported in part by the WSU Monoclonal Antibody Center.

